# Intrinsic structure of lipoplexes embedded in polyelectrolyte multilayers

**DOI:** 10.1101/2025.10.10.681465

**Authors:** Maria Krabbes, Vincent Kampik, Mathilde Büttner, Leonard Kaysser, Emanuel Schneck, Chen Shen, Christian Wölk

## Abstract

The functionalization of surfaces with therapeutically applicable nucleic acid carriers provides promising strategies in biomedical research to develop therapies which focus on local nucleic acid delivery. One such approach is the embedding of lipoplexes (LPXs) in polysaccharide-based polyelectrolyte multilayers (PEMs). PEMs based on hyaluronic acid and chitosan lead to efficient embedding of customized LPX connected with good biological activity. However, although quantitative evaluation demonstrates LPX embedding, information on the embedded LPXs has been missing. In this study we used synchrotron-based grazing-incidence small angle x-ray scattering to investigate the effects of the change in the chemical environment caused by the embedding into PEMs on the LPX’s internal mesoscopic structure. While the lamellar character of the LPXs was preserved, the repeat distance was affected by the embedding into polysaccharide-based coatings.

## 1. Introduction

During the last decade, lipid-based nanoparticle systems evolved to an effective class of carriers to deliver therapeutic nucleic acids into cells because of beneficial characteristics regarding e.g. safety, tolerability, ability to re-dose, transfer capacity, and ability to apply structural design-concepts (Cullis and Felgner, 2024). Lipid nanoparticles (LNPs) and lipoplexes (LPXs) are the most prominent lipid-based delivery systems for nucleic acids. Commonly, the nanocarriers can be applied systemically via i.v. injection or locally (e.g. in muscular tissue for vaccination) (Cullis and Felgner, 2024). Hence, biomedical applications of nucleic acid therapeutics can benefit from a nanoparticle reservoir for local release. A promising strategy is the application of hydrogels or solid scaffolds loaded with nucleic acid carriers as gene-activated matrices in the field of regenerative medicine for e.g. cartilage or wound regeneration (Wang et al., 2024) (Palomeque Chávez et al., 2025). Special cases are gene-activated surface coatings based on polyelectrolyte multilayers (PEMs) with embedded LPXs with the aim do design substrate-mediated gene delivery, while currently PEMs composed of at least one polysaccharide component are used: Liu et al. designed a PEM coating based on hyaluronic acid and LPXs (Liu et al., 2011); Holmes et al. proposed a coating strategy based on hyaluronic acid, chitosan, and LPX (Holmes and Tabrizian, 2013); a minimalistic coating of polyallylamine, chondroitin sulfate, poly-L-lysine, and LPX was designed by Carvalho et al. (Carvalho et al., 2022); and from our laboratory, Husteden et al. investigated two different systems embedding LPXs, either in chondroitin sulfate/collagen PEMs (Husteden et al., 2023), or in hyaluronic acid/chitosan PEMs (Husteden et al., 2020). Although the binding and embedding of LPXs in the PEM systems was proven and also quantification approaches were presented, the effect of the embedding process on the internal structure of the LPXs was never studied.

In the present work, we investigate the effect of the interaction of LPX with the polyelectrolytes of the PEMs on the mesoscopic LPX structure. The key technique was grazing incidence small angle x-ray scattering (GISAXS), a technique that allows to determine the LPX mesostructure on ultra-thin films like PEM coatings. As test system, PEMs composed of hyaluronic acid (HA) and chitosan (CHI) were chosen (see Figure 1), because the preparation conditions for these materials were intensively studied in our group (Husteden et al., 2020) (Krabbes et al., 2025a). Also the embedded LPX, composed of the ionisable lipid OH4 (designed in our group) and the co-lipid DOPE (structures of both lipids are given in Figure 1), have already been characterized physicochemically as nanoparticles in dispersion (Janich et al., 2016). Our findings demonstrate that the embedding of LPXs in the PEMs can affect their structure.

**Figure 1.**
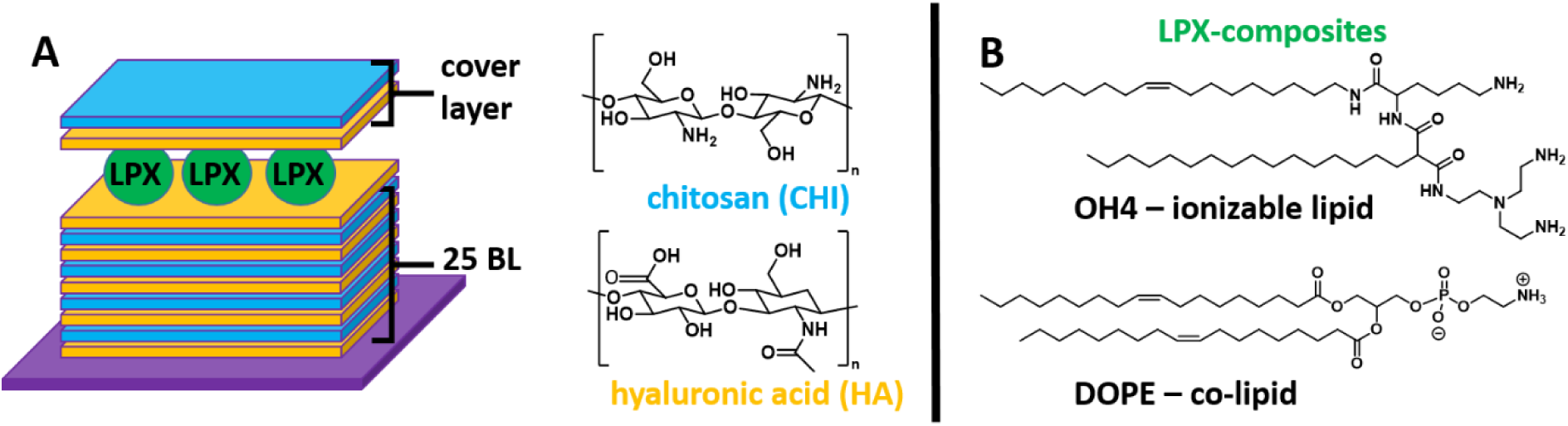
(A) Schematic illustration of the LPX-functionalized PEM system. The PEM components are chitosan (CHI, polycation) and hyaluronic acid (HA, polyanion). The surface coating is built of a base-layer comprising 25 HA/CHI bilayers (BL) and a final HA layer to provide a negative charge for LPX adsorption. A cover layer composed of HA and CHI terminates the LPX-loaded PEM coating. (B) Lipid structures of the LPX components, OH4 and DOPE.

## 2. Materials and Methods

### 2.1 Materials

If not stated otherwise, the chemicals were acquired from Merck KGaA (Darmstadt, Germany). The phospholipid 1,2-di-(*9Z*-octadecenoyl)-*sn*-glycero-3-phosphoethanolamine (DOPE) was from Avanti Polar Lipids (Alabaster, USA). The ionizable lipid *N*-{6-amino-1-[*N*-(9*Z*)-octadec-9-enyl-amino]-1-oxohexan-(2S)-2-yl}-*N′*-{2-[*N,N*-bis(2-aminoethyl)amino]ethyl}-2-hexadecylpropandiamide (OH4) was synthesized in our research group (Janich et al., 2014). The plasmid DNA (pDNA, 3.5 kbp, 1 mg/mL stock, product abbreviation pCMV-GFP), encoding for the green fluorescent protein, was purchased from PlasmidFactory (Bielefeld, Germany).

### 2.2 Polyelectrolyte Solutions

Sodium acetate buffer at pH 5.5 was prepared by combining a 0.2 M sodium acetate solution (Merck, Darmstadt, Germany) and glacial acetic acid (AppliChem, Darmstadt, Germany) and an adjustment with deionised water to a final concentration of 0.1 M acetate. The polyelectrolytes sodium hyaluronate (HA, MW ≈ 1.3 MDa; Kraeber & Co GmbH, Ellerbek, Germany) and chitosan 85/500 (CHI, MW ≈ 0.2-0.4 MDa, degree of deacylation ≈ 85 %; Heppe Medical Chitosan GmbH, Halle (Saale), Germany) were separately dissolved in the 0.1 M acetate bufferto a concentration of 2 mg/mL by stirring overnight and subsequent filtration using Minisart^®^ high flow syringe filters (4,4′-Dichlorodiphenyl sulfone-4,4′-dihydroxydiphenyl sulfone copolymer, pore size 0.45 µm; Sartorius, Göttingen, Germany).

### 2.3 Base-Layer PEM Preparation

Silicon wafers (10.0 mm × 15.0 mm, 0.6 mm thick, (100) face, Si-Mat, Kaufering, Germany) were used as substrates for the PEM base-layer preparation for characterization with X-ray reflectivity (XRR) and GISAXS. The wafers were cleaned with an RCA-1 protocol (https://doi.org/10.1021/acsami.9b18968) and kept in Milli-Q water for no more than 6 days. For the determination of the loading efficiency using a gel electrophoresis-based quantification, glass cover slips with a diameter of 13 mm and a thickness of 0.13–0.16 mm (Karl Hecht GmbH & Co KG, Sondheim vor der Rhön, Germany) were used as substrates. For the PEM coating of the substrates, a DR 0 Layer-by-Layer deposition robot (Riegler & Kirstein GmbH, Potsdam, Germany) controlled by Dipp3dWin software was used allowing for an automated PEM formation with established dipping protocols (Krabbes et al., 2025a). To fix the wafers or cover slips for the dipping process, custom designed substrate holders were utilized (Krabbes et al., 2025a). Briefly, PEM preparation was performed by alternating incubation of the substrates in the polyelectrolyte solutions following the sequence: 5 min HA solution incubation, 2.5 min washing step in 0.1 M acetate buffer pH 5.5, 5 min CHI solution incubation, 2.5 min washing step in 0.1 M acetate buffer pH 5.5. This sequence was repeated until the base-layer PEM [HA, CHI]_25_HA was completed. The pre-formed base-layer PEMs were stored separately in a 12-well plate in 0.1 M acetate buffer pH 5.5 at 4°C until used for measurements or further processing. The process is schematically illustrated in Figure 2A.

**Figure 2.**
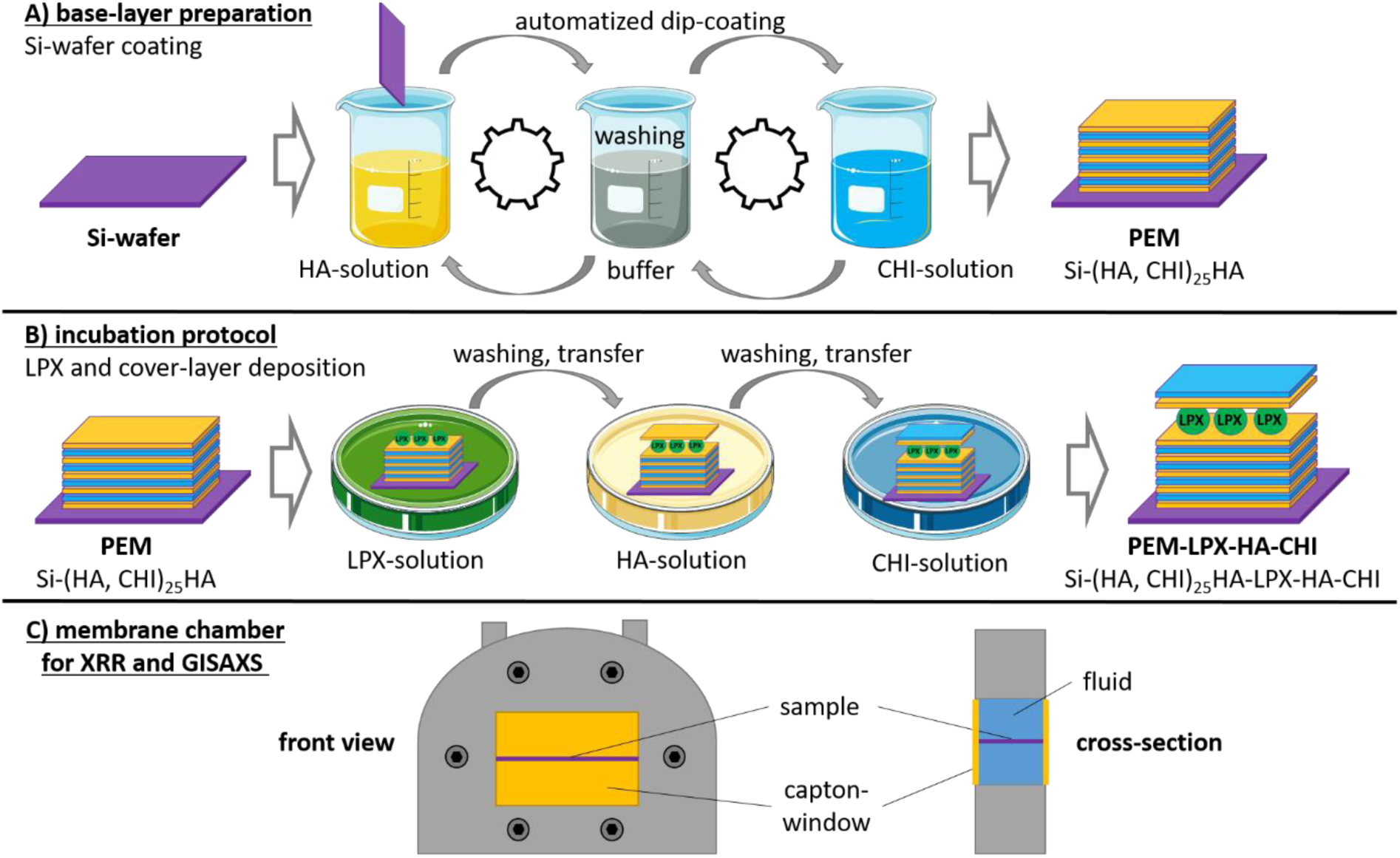
Illustration of the coating process applied to the silicon wafers (panel A). The base-layer PEM (sequence: Si-(HA, CHI)_25_HA was prepared with an automated dip coating protocol. The LPX deposition and cover-layer adsorption was performed by an incubation protocol in 12-well plates (panel B). Panel C displays the polyether ether ketone (PEEK) chamber body for placing the coated silicon wafer in aqueous environment, with the Kapton foil windows, used for synchrotron x-ray experiments. The two Kapton windows were clamped by two Aluminium window frames (not shown) from both sides of the chamber body. The frames consisted of water channels for temperature control. A Pt100 temperature sensor (not shown) was embedded into the PEEK body in contact with the aqueous solution.

### 2.4 Liposome Preparation

OH4 and DOPE were separately dissolved in a chloroform/methanol mix (8/2 volume/volume) at a concentration of 2 mg/mL and mixed to a final molar ratio of 1:1. The organic solvent was then removed using a rotary evaporator at 500 mbar for 30 min followed by further solvent evaporation at 10 mbar for 1.5 h. The dried lipid film was then re-solvated in 0.1 M acetate buffer pH 5.5 by shaking at 600 rpm for 30 min at 55°C using an Eppendorf ThermoMixer C (Eppendorf SE, Hamburg, Germany). To form liposomes, sonication at 55°C and 37 kHz for 15 min using an Elmasonic P sonication bath (Elma Schmidbauer GmbH, Singen, Germany) was performed. The final total lipid concentration in the liposome dispersion was 1 mg/mL. Liposomes were stored at 4°C until use and sonication at room temperature was conducted for 5 min right before LPX preparation.

### 2.5. Lipoplex Preparation

For the preparation of LPX, the pDNA solution was added to the cationic liposomes in one step at an N/P ratio of 4 (ratio of primary amines of OH4 - N - to phosphate groups of the nucleic acid - P) in 0.1 M sodium acetate buffer (pH 5.5). The LPXs were prepared under gentle mixing by pipetting and afterwards shaken for 15 min at 300 rpm with a ThermoMixer C (Eppendorf SE, Hamburg, Germany).

### 2.6. Lipoplex Deposition on Base-layer PEMs - Incubation protocol

LPXs were diluted to “LPX-loading dispersions” of the following concentrations: 4.33 ng_(DNA)_/µL (1x c(LPX)), 8.66 ng_(DNA)_/µL (2x c(LPX)), 17.32 ng_(DNA)_/µL (4x c(LPX)) and 34.64 ng_(DNA)_/µL (8x c(LPX)). For LPX loading of base-layer-PEMs an incubation protocol was used (see schematic illustration in Figure 2B). The storage buffer of the PEMs was removed from the wells and 1050 µL the appropriate LPX loading dispersion were pipetted to each base-layer PEM in the 12-well plate and incubated for 2 h on the 3D rocker shaker (VWR International GmbH, Darmstadt, Germany) at speed/tilt 8. After LPX incubation, PEMs were washed twice with buffer for 5 min. If a deposition of HA and CHI cover layers was applied, the PEMs were subsequently incubated for 10 min with 1050 µL HA solution, washed twice with buffer, and were then incubated for 10 min with 1050 µL CHI solution and finally washed twice with buffer. All steps were conducted on the 3D rocker shaker.

### 2.7 Synchrotron X-ray Experiments

GISAXS and XRR measurements were used to examine the change of the LPX mesostructure and the structure of the PEM coating upon the embedding process. The silicon wafers, either PEM-coated or non-coated (depending on the experiment), were mounted into the SDU-Odense membrane chamber (AG. Klösgen, University of Southern Denmark, Denmark) (Figure 2C). The wafers were inserted centrally and, horizontally, and tightly sealed in the chamber. Through a side port, 1.5 mL of the acetate buffer was filled into the compartment for the measurements buffer (≈500 µL below and ≈ 1000 µL above the wafer). The chamber was mounted in the experimental setup and then thermostatically adjusted to 22.0±0.2 °C for about 10 min prior to the X-ray measurement. The real-time temperature of the aqueous phase was monitored over the preparation and measurement by an integrated Pt100 sensor in the chamber.

For most of the measurements, the PEM coating (with or without LPX embedding) was performed *ex-situ*, before mounting the silicon wafer in the chamber. For the *in-situ* evaluation of LPX deposition on the silicon wafers with base-layer PEM coating or empty silicon wafers the LPX deposition was performed in the liquid chamber. 500 µL of the buffer were removed from the chamber and replaced by 500 µL of a LPX dispersion at 33.3 ng_(DNA)_/µL, resulting in a nominal concentration of ≈ 17 ng_(DNA)_/µL. During this loading process, the wafer was always immersed in the buffer.

The measurements were conducted on a Kohzu 6-circle diffractometer at the beamline P08 of the PETRA III synchrotron (DESY, Hamburg Germany) (Seeck et al., 2012). An incident beam at 18keV with focus mode was used, providing a beam size of 0.07 mm x 0.3 mm (vertical x horizontal). A set of guard slits and a pre-sample pinhole with a diameter of 0.8 mm were used for reducing the beam path scattering background before the sample. The XRR signal was measured with a Pilatus 100k (Dectris, Switzerland) at 1000 m from the center of the sample surface, positioned at two times the sample incident angle with respect to the incident beam (θ-2θ geometry). The reflected intensity was integrated in a detector area of 1.0 mm x 1.5 mm (vertical x horizontal), and normalised to the incident beam intensity. The GISAXS signal was measured with an Eiger2X 1M detector (Dectris, Switzerland) placed at 1218 mm from the sample surface center. The direct and the reflected beam were blocked by a 1.2 mm wide Tungsten finger beamstop placed at ≈300 mm from the sample. The angle of incidence with respect to the sample plane was set as a series of 0.07°, 0.2° and 0.5°. Note that the lowest angle corresponds to a total-reflection condition of the silicon surface (critical angle: ∼0.095°). The buffer GISAXS background signal of the chamber setup was measured by lowering the sample by 1 mm, to be subtracted from the sample surface scattering signal. Azimuthal integration of the GISAXS data was performed using pyFAI library (Ashiotis et al., 2015) to obtain one-dimensional (1D) GISAXS diffractogram for further analysis.

### 2.8 Small Angle X-ray Scattering

For SAXS measurements of LPXs dispersions, three different sample preparations were performed (named sample 1 to 3). The liposome and LPX protocols had to be adapted to achieve LPX preparations of concentrations sufficient for SAXS measurments. The dry lipid films for liposome preparation were prepared from lipid stocks of 10 mg/mL. Liposomes for LPX sample 1 and 3 were prepared by adding 100 mM acetate buffer pH 5.5 to the lipid film to yield a 10.6 mg/mL lipid dispersion which was treated by 10 min sonication at 50°C. To prepare LPX sample 1, 200 µL liposome dispersion (10.6 mg_(total lipid)_/mL) was added to 330 µL pDNA solution (1 mg/mL in solvent from supplier) in one step and incubated for 20 min at 22°C and 120 rpm for LPX formation. The turbid LPX dispersion was centrifuged for 30 min at 20 °C and 30,000 g. The supernatant was removed and the LPX pellet re-suspended in 50 µL 100 mM acetate buffer pH 5.5 to yield a final LPX concentration of 6.6 mg_(DNA)_/mL. To prepare LPX sample 3, 50 µL pDNA solution (6.6 mg/mL in PCR grade water) were added to 200 µL liposome dispersion (10.6 mg_(total lipid)_/mL) in one step and incubated for 20 min at 22°C and 120 rpm for LPX formation. The turbid LPX dispersion had a final concentration of 1.32 mg_(DNA)_/mL. For the preparation of the LPX sample 2, A dried lipid film of 2.1 mg total lipid was dispersed in 50 µL 100 mM acetate buffer pH 5.5 (10 min sonication at 50°C). 50 µL pDNA solution (6.6 mg/mL in PCR grade water) were added to the lipid dispersion followed by incubated for 20 min at 22°C and 120 rpm. The LPX concentration was 3.3 mg_(DNA)_/mL. All samples were transferred in borosilicate glass capillaries (1.5 mm outside diameter, 0.01 mm wall thickness; WJM-Glas / Müller GmbH, Berlin, Germany). The glass capillaries were sealed for measurements.

The SAXS measurements were carried out at the Biomaterials Department of Max Planck Institute of Colloids and Interfaces (MPIKG, Potsdam, Germany) with a Bruker Nanostar 2 (Bruker, Billerica, MA, USA), equipped with a 2D Vantec-2000 detector and a microfocus X-ray source (IµS) with 1.542 Å wavelength (Cu Kα) and a focal spot size of 115 µm. The following measurement parameters were set: 50 kV voltage, 600 µA current, 1071 mm sample-detector distance. As calibration material, silver behenate was used. The capillaries were placed in a sample holder and the total scanning time was 85 h. For data processing SAXS: Small Angle X-ray Scattering System V4.1.61 software (Bruker, Billerica, MA, USA). As background measurements, capillaries filled with buffer, empty capillaries, and the empty sample holders were measured.

### 2.9 Quantification of the DNA loading

The pDNA loading of base-layer PEMs was quantified indirectly by determination of the pDNA concentration in the supernatant of the LPX loading dispersion and washing solutions. The supernatant and washing solutions were collected after PEM incubation with the LPX loading dispersion. The samples of the washing solutions were combined for analysis. For total pDNA quantification a disintegration/decopmlexation of LPX was necessary. For this purpose, 150 µL of the collected solutions were transferred into micro-vessels and mixed with a 60 mg/mL heparin solution (sodium heparin from porcine intestinal mucosa, ≥180 USP units/mg) (Sigma-Aldrich, Taufkirchen, Germany) in Dulbecco’s PBS (AppliChem, Darmstadt, Germany) to a final concentration of 2 μg/μL heparin in the samples. The samples were shaken for 5 min at 1500 rpm in a ThermoMixer C (Eppendorf SE, Hamburg, Germany). Subsequently, an equal volume of isopropanol (solvent temperature 0°C) was added to the samples and incubated for 20 min at -20°C to precipitate the pDNA. After centrifugation at 21,300g for 10 min at 4°C, the solvent was carefully removed from the samples. 500 µL of ethanol (solvent temperature 0°C) was used to gently wash the resulting pellet. After removal of the ethanol, the samples were resuspended in 20 µL water (pH 8). To quantify the pDNA amount, an additional digestion was implemented to linearize the plasmid, due to variations in the content of supercoiled and open circular pDNA. A volume of10 µL of the DNA solution were separately mixed with 7,6 µL water, 2 µL of rCutSmart Buffer™ and 0,4 µL XhoI ™ (both from New England Biolabs, Ipswich, MA, USA) restriction endonucleases and the preparation was incubated for 1 h at 37 °C. As a standard for quantification pDNA was treated the same way. Furthermore, TriTrack DNA Loading Dye (6X) (Thermo Fisher Scientific, Darmstadt, Germany) was added, and the samples were applied to agarose gels. The gels were prepared in a concentration of 1 % agarose in TAE buffer with thiazol-orange. The GeneRuler 1 kb Plus DNA ladder (Thermo Fisher Scientific, Darmstadt, Germany) and pDNA in different concentrations for the calibration curve for assessment were also loaded on the gel. The electrophoresis was performed for 1 h at 120 V and the gels were evaluated using a GelDoc Go Imaging System (Bio-Rad Laboratories, Hercules, USA). The gel is shown in the Supporting Information.

### 2.10 Determination of the Lamellar Repeat Distance

The lamellar repeat distance *d* was calculated from the peak maximum of a Bragg reflection as *d* = (*n* · 2π)/*q*, where *q* is the magnitude of the scattering vector and *n* is the order of the Bragg reflection. The calculated *d* values are given as mean for 3 different positions in the capillary (SAXS) or 4 different positions on the silicon wafer (GISAXS). For SAXS and GISAXS, a Lorentz fit was used to determine the peak maximum.

## 3. Results and Discussion

### 3.1 SAXS measurements

SAXS measurements were performed to identify the mesophase substructure of the LPX nanoparticles composed of the lipid composite OH4/DOPE encapsulating pDNA at a NP (primary amine to phosphate) loading ratio of 4 (OH4/DOPE NP4 LPX) in acetate buffer pH 5.5 (100 mM), the processing buffer used for embedding of LPX in polysaccharide PEMs (Krabbes et al., 2025a). Earlier studies of OH4/DOPE NP4 LPX in MES buffer pH 6.5 (100 mM) was reported to result in a lamellar L_α_^c^ LPX structure (Figure 3B), a supermolecular multilamellar mesophase structure with alternating lipid bilayer and DNA monolayers (Rädler et al., 1997) with lamellar repeat distance *d* = 69 Å (Janich et al., 2016). For OH4/DOPE NP4 LPX in acetate buffer pH 5.5 (100 mM) the Bragg peaks also indicated a lamellar L_α_^c^ LPX structure showing the first and second order peak, at *q*_(001)_ and *q*_(002)_ = 2 *q*_(001)_, respectively (see Figure 3A and Table 1). The determined *d* value (see schematic illustration in Figure 3B) ranged between 64.2 and 66.6 Å in dependence of the preparation procedure of sample 1-3 (three different procedures were chosen to achieve LPX preparations with high concentrations). Consequently, a significantly lower *d* value was observed compared to the LPX in MES buffer pH 6.5. The experimental determination of the apparent pKa (pKa_app_) value of homologue OH4 derivatives via determination of lipid charge state by counterion quantification, using total reflection x ray fluorescence measurements at the air/water interface on lipid monolayers with bromide as counterion, indicate a pKa_app_ ≈ 6 (Tassler et al., 2017). Although recent research demonstrated, that the protonation degree can differ significantly from the value expected from the Henderson– Hasselbalch equation (Grava et al., 2023), we want to use the above mentioned pH for theoretical consideration. Calculating the ratio of RNH_3_^+^/RNH_2_ using the Henderson-Hasselbalch equation [pH = pKa-lg(c{R-NH_3_^+^}/ c{R-NH_2_})] and a pKa of 6 results in a R-NH_3_^+^/R-NH_2_ ratio of 0.32 for MES pH 6.5 and 3.16 for acetate pH 5.5. Consequently, the charge density of the lipid formulation is higher in the acetate buffer. An increase of positive charge density of the lipid formulation caused by a higher protonation degree can cause a decrease in the *d* value of L_α_^c^ LPX according to described investigations in literature (Hubčík et al., 2014) (Uebbing et al., 2020) (Wilhelmy et al., 2025).

**Figure 3.**
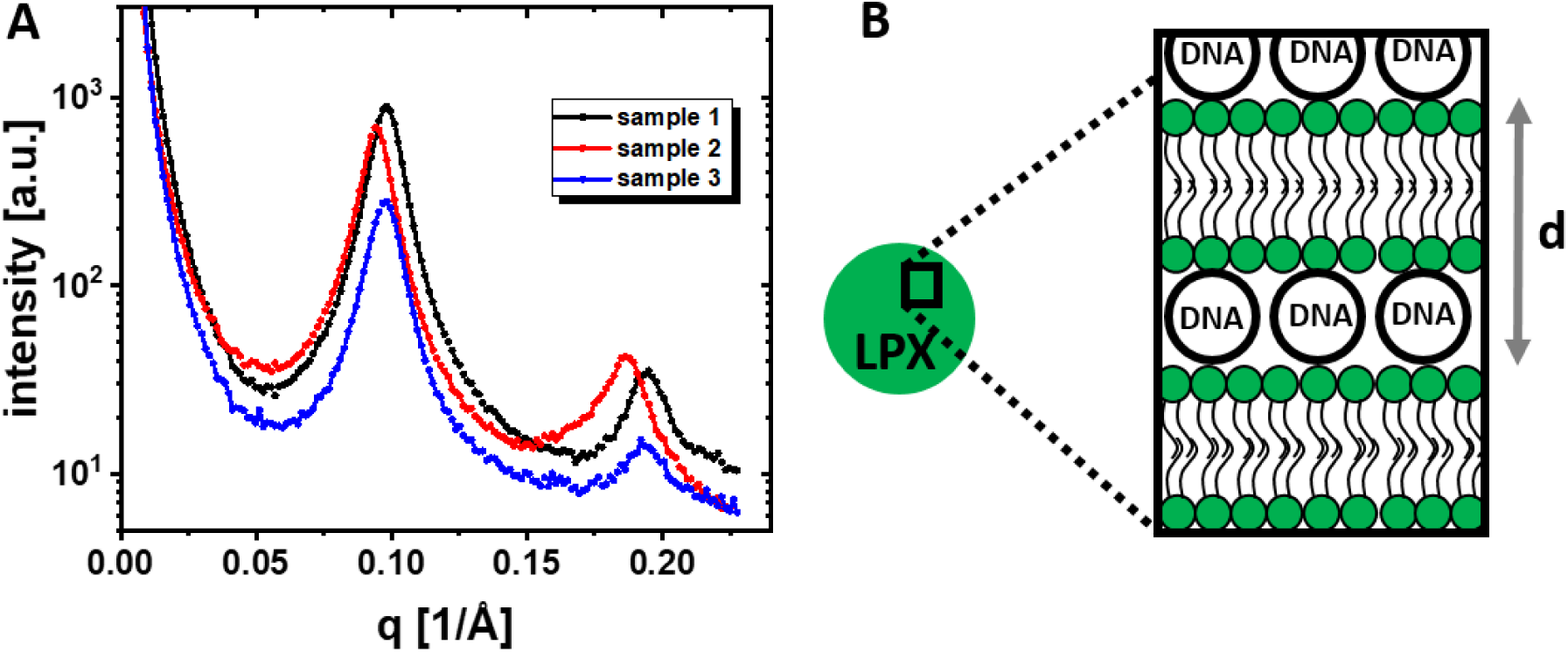
A) SAXS pattern of OH4/DOPE N/P4 LPX in acetate buffer pH 5.5. The three different samples were prepared by different protocols and resulted in different final LPX concentrations (see Table 1). B) Illustration of the L_α_^c^ LPX structure

**Table 1.** Summary of SAXS results of OH4/DOPE NP4 LPX in acetate buffer pH 5.5 for the three different samples.

|  | sample 1 | Sample 2 | Sample 3 |
| --- | --- | --- | --- |
| LPX concentration | 6.6 mg<sub>(DNA)</sub>/mL | 1.32 mg<sub>(DNA)</sub>/mL | 3.3 mg<sub>(DNA)</sub>/mL |
| $q_{(001)}^*$ | 0.098 1/Å | 0.094 1/Å | 0.097 1/Å |
| $q_{(002)}^*$ | 0.194 1/Å | 0.186 1/Å | 0.193 1/Å |
| $q_{(002)}$ calculated as<br>$2 \times q_{(001)}$ | 0.196 1/Å | 0.189 1/Å | 0.195 1/Å |
| $d_{\text{lamellar}}^{\#}$ | 64.2 Å | 66.6 Å | 64.5 Å |

\* mean of Lorenz-fits of the 3 different curves, given in the supplemental information Figures S1-S3)

### 3.2. Characterization of LPX Embedded in Polysaccharide PEMs

For LPX embedded in PEMs composed of HA and CHI, the chemical environment of the lipid nanoparticles changes drastically compared to the dispersion of the nanoparticles in aqueous medium. Hence, LPX supermolecular assembly is based on electrostatic interaction and entropic effects (Meidan et al., 2000) (Pozharski and MacDonald, 2003). It is thus plausible to expect structural changes of the LPXs caused by the embedding, and the characterization of such changes is the central aspect of the present work.

We performed XRR measurements to check whether or not the PEM coatings have a layered internal structure themselves, as described in literature (Lvov et al., 1993), or if the nanoparticles cause structural features which can be detected by XRR. The reflectivity measurements indicate no relevant electron density modulation within the PEMs (for more detailed discussion see supporting information, section 6 and Figure S17). According to earlier reports, HA/CHI multilayers are characterized by an exponential growth of loosely associated polyelectrolyte layers, by soft and hydrated structures, and the tendency of CHI to diffuse in the PEM (Kujawa et al., 2005) (Richert et al., 2004) (Almodóvar et al., 2011).

GISAXS was the second technique to screen the surface coating for structural features. The azimuthally integrated GISAXS data of the functionalized base PEM (sequence on the silicon wafer: silicon-(HA, CHI)_25_HA, Figure 4D/E, curve labelled PEM) show a curve with monotonic intensity decay and the absence of Bragg reflection peaks, which is consistent with the XRR result. The deposition of LPX using the standard LPX loading dispersion and addition of the cover layer (sequence on the silicon wafer: Si-(HA, CHI)_25_HA-LPX-HA-CHI) changes the diffractogramm (Figure 4D/E, curve labelled 1 x c(LPX)). The system with embedded LPX exhibits a single Bragg peak which is the first order reflection (indicated as *q_(001)_*) of the L_α_^c^ LPX structure. Compared to the SAXS results for LPX in dispersion, the peak position is shifted to higher q values, which indicates that the embedding into PEM decreases the *d* spacing in LPX by 7 to 9 Å to d ≈ 57 Å (compare Table 1 with lamellar structure 2 in Table 2). Hence the question occurs, if it is possible to embed higher amounts of LPX into the PEM structure. Consequently, the concentration of the LPX loading dispersion in the LPX-deposition step was increased by factors 2, 4, and 8. Initially it was planned to terminate all the coatings with a cover-layer to the final sequence si-(HA,CHI)_25_HA-LPX-HA-CHI. But, due to a visible aggregation of LPX on the PEMs incubated with the 8 x c(LPX) dispersion, a cover-layer deposition was avoided and the sample was only used for GISAXS measurements and not for quantification of LPX loading. The observed aggregation also resulted in an inhomogeneous distribution of LPX on the PEM, as demonstrated by pronounced variations of the signal intensities in the GISAXS position scan (Supporting Information Figure S15). Evaluating the effectivity of LPX embedding in this set of experiments, the LPX loading given as amount DNA normalized to the surface was significantly increased with the increase of the concentration of the loading in the screened range (Figure 4B). Hence, the possibility to incorporate higher amounts of LPX into the PEM was demonstrated. The diffractogram also changes with increasing LPX mass deposited (Figure 4D/E). With increasing amount of LPX loading, the peak signal increases in intensity. The area of the signal is proportional to the volume fraction of the LPX in the measurement area of the sample in the GISAXS setup. Consequently, a higher peak area corresponds a higher amount of embedded LPX. The second observation was a clear appearance of two different first order peaks of the lamellar LPX structure for the 2 x c(LPX) and 4 x c(LPX) sample (see reflex labelled L1 and L2 in Figure 4D/E and Table 2). The reflex at ≈0.1 1/Å is comparable to the L_α_^c^ LPX structure of the LPX in solution determined by SAXS, and the reflex at ≈0.11 1/Å is the structure which was observed for the lowest investigated amount of embedded LPX which was applied to the samples (1 x c(LPX)). Also, in the sample with the highest loading concentration investigated (8 x c(LPX)), the sample with visible LPX aggregation which was prepared without cover layer, both reflexes were visible (Figure 4F). Additionally, samples with the 1 x c(LPX) loading concentration were prepared with HA or HA/CHI cover layer coating of LPX and stored for 4-5 days in buffer. Also here the reflex of the L2 LPX structure was observed (Supporting Information Figure S7 Table S1), what is in line with the results presented in Figure 4D/E for the 1 x c(LPX) sample.

**Figure 4.**
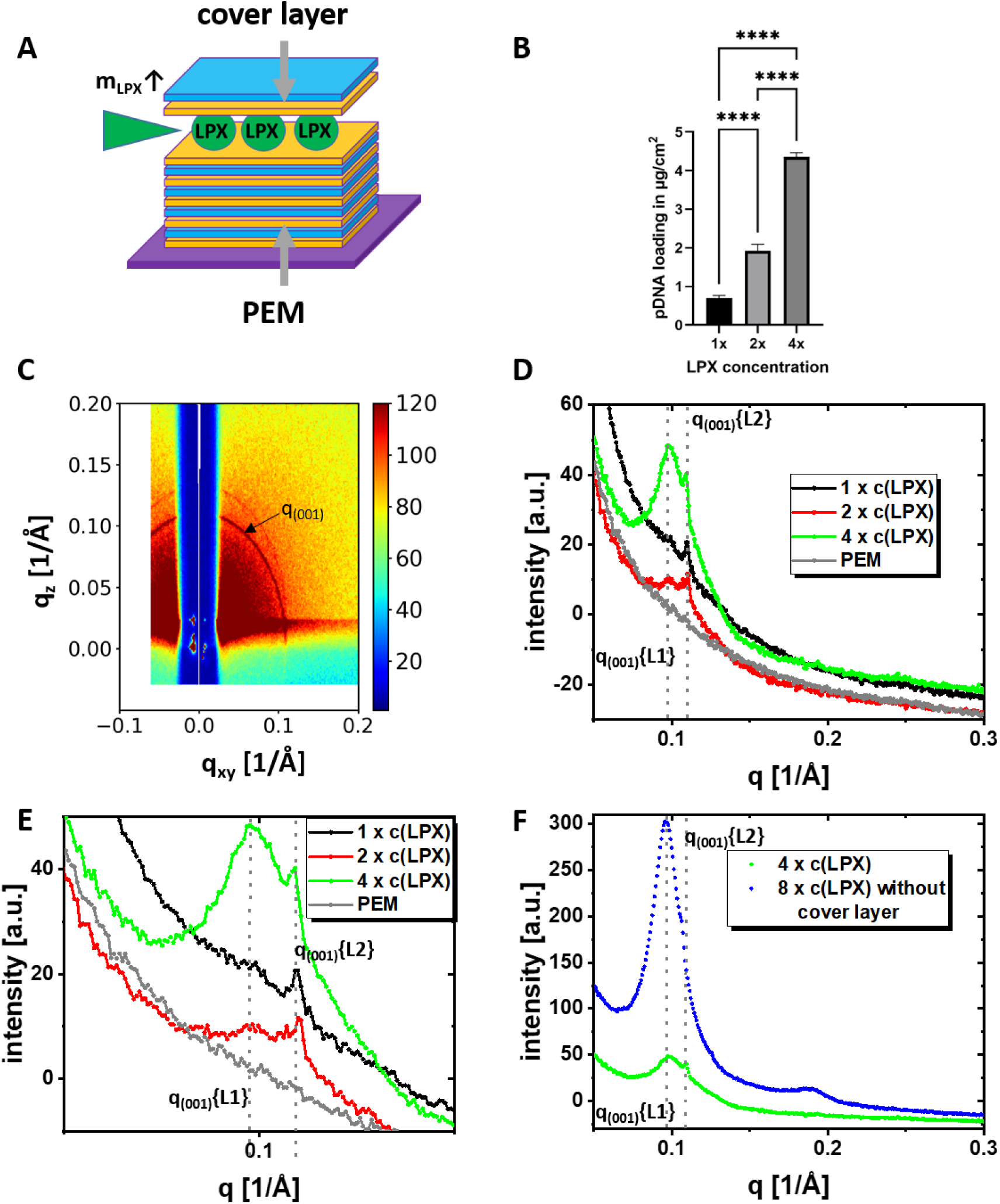
A) Schematic illustration of the investigated LPX-loaded PEMs. The mass of embedded LPX was increased by performing the LPX deposition with increased LPX concentration (c(LPX)) in the LPX loading dispersions. B) Determination of loading efficiency of PEM-films with LPX cargo by indirect DNA quantification using gel electrophoresis given in absolute DNA amount in g/cm^2^ for PEMs loaded with the LPX loading dispersion single, double and quadruple concentration (labelled with 1x, 2x, and 4x). Given values are mean ± standard deviation of n = 3. One way anova with Tukey post hoc test. Was performed to check for statistical significance (**** = p < 0.0001). C-F) GISAXS experiments of si-(HA, CHI)_25_HA-LPX-HA-CHI samples using different LPX concentrations in the LPX deposition step. C) example 2D GISAXS pattern before azimuthal integration. The first order peak of the L_α_^c^ LPX structure is labelled as q_(001)_, and has the form of a Scherrer ring. D) Azimuthally-integrated 1D GISAXS diffractogram of PEMs with different LPX loading concentrations. The q_(001)_ reflexes of two different lamellar phases were observed, labelled with q_(001)_ {L1} and q_(001)_ {L2}. The concentration of the LPX dispersion for the LPX loading were varied from the standard concentration (1 x c(LPX) to higher concentrations, namely 2 x, and 4 x (representing the factor of concentration increase). E) Detailed q range from (D). F) GISAXS experiments of si-(HA,CHI)_25_HA-LPX-HA-CHI with 4 x c(LPX) and si-(HA, CHI)_25_HA-LPX with 8 x c(LPX) loading. The GISAXS diffractograms of the samples presented in (D), (E), and (F) are shown at 4 different positions on the sample in the supporting information (Figure S12-15) to demonstrate homogeneity of the samples.

**Table 2.** LPX Bragg peak position obtained from GISAXS measurement of OH4/DOPE NP4 LPX on or embedded in PEMs as mean of 4 different positions.

| | | lamellar<br>structure 1<br>$q_{(001)}\{L1\}$<br>[d] | lamellar<br>structure 2<br>$q_{(001)}\{L2\}$<br>[d] | LPX loading<br>dispersions:<br>concentration<br>[volume] | coverlayer<br>on top of<br>LPX |
| --- | --- | --- | --- | --- | --- |
| *ex-situ* sample |  |  |  |  |  |
| 1 x c(LPX) |  | - | 0.1096 1/Å<br>[57.3 Å] | 4.33 ng<sub>(DNA)</sub>/μL<br>[1050 μL] | yes |
| 2 x c(LPX) |  | 0.1016 1/Å<br>[61.8 Å] | 0.11 1/Å<br>[57.1 Å] | 8.66 ng<sub>(DNA)</sub>/μL<br>[1050 μL] | yes |
| 4 x c(LPX) |  | 0.0992 1/Å<br>[63.4 Å] | 0.1092 1/Å<br>[57.6 Å] | 17.32 ng<sub>(DNA)</sub>/μL<br>[1050 μL] | yes |
| 8 x c(LPX) |  | 0.0961 1/Å<br>[65.4 Å] | 0.1063 1/Å<br>[59.1 Å] | 34.64 ng<sub>(DNA)</sub>/μL<br>[1050 μL] | no |
| *in-situ* sample |  |  |  |  |  |
| 4 x c(LPX) | 45<br>min | 0.0987 1/Å<br>[63.6 Å] | 0.1065 1/Å<br>[59 Å] | *in-situ* deposition<br>starting concentration<br>17 ng<sub>(DNA)</sub>/μL<br>No coverlayer |  |
|  | 60<br>min | 0.0962 1/Å<br>[65.3 Å] | 0.1059 1/Å<br>[59.4 Å] |  |  |
|  | 90<br>min | 0.0977 1/Å<br>[64.3 Å] | 0.1065 1/Å<br>[59.0 Å] |  |  |
|  | 120<br>min | 0.0972 1/Å<br>[64.7 Å] | 0.1066 1/Å<br>[59 Å] |  |  |
|  | 180<br>min | 0.0972 1/Å<br>[64.6 Å] | 0.1065 1/Å<br>[59 Å] |  |  |
|  | 240<br>min | 0.0969 1/Å<br>[64.9 Å] | 0.1066 1/Å<br>[58.9 Å] |  |  |
|  | 540<br>min | 0.0956 1/Å<br>[65.7 Å] | 0.1062 1/Å<br>[59.2 Å] |  |  |

After observing two different lamellar LPX structures, the process of LPX deposition was screened *in-situ* (see Figure 5) to get more insights in the process of deposition. The concentration of LPX in the chamber was comparable to the 4 x c(LPX) experiment mentioned above. Within the screened time frame of 540 min, the intensity of the LPX derived diffraction pattern increased monotonically with time. As mentioned before, the intensity is correlated to the volume fraction of LPX and consequently the adsorbed amount of LPX. Hence, the adsorption of LPX on the PEM was efficient and it is possible to increase the deposited amount of LPX by the increase of incubation time. The driving force of the adsorption can be attributed to the electrostatic interaction of the LPX (positive zeta potential (Krabbes et al., 2025a) (Krabbes et al., 2025b) with the negatively charged HA of the basal PEM. Further, the *in-situ* experiment revealed the coexistence of the two structural LPX types (L1 and L2). Obviously, the adsorption of the cover layer is not necessary to convert L1 LPX to L2 LPX, which is supported also by the observation of both LPX phases for 8 x c(LPX) experiment mentioned above. Nevertheless, the question arises if the effect of the appearing L2 LPX structure is generally caused by the LPX adsorption on surfaces. Hence, we performed an *in-situ* experiment of the LPX adsorption to the bare silicon wafer for 2 h (Figure 5D/E). Also here a certain affinity of the LPX to the surface was observed by detecting Bragg reflections. The signal intensity was less compared to the *in-situ* adsorption to PEM coated silicon for the same adsorption time, indicating the benefit of the HA LPX interaction for the adsorption process. Further, no signal of L2 LPX was observed, indicating that the contact to the PEM is needed for the transition.

**Figure 5.**
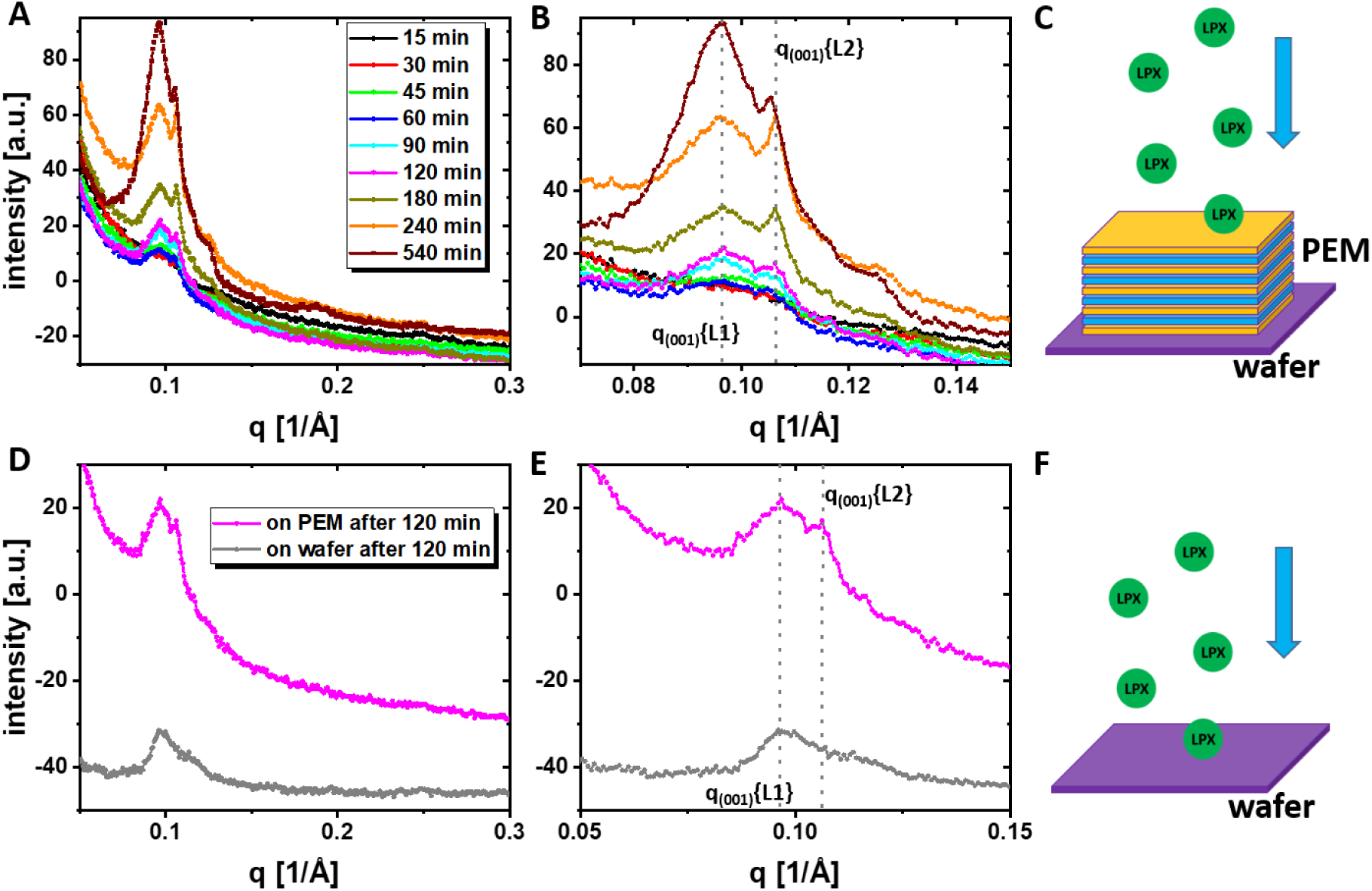
GISAXS *in-situ* LPX deposition experiments of PEMs Si-(HA, CHI)_25_HA or unmodified silicon wafers with LPX in the supernatant for sedimentation. In the liquid above the modified or unmodified silicon wafer the LPX concentration was 17 ng_(DNA)_/µL. A) 1D GISAXS diffractogram of si-(HA, CHI)_25_HA PEMs adsorbing LPX from solution after different time points in the LPX containing medium. The two different lamellar phases were labelled with q_(001)_ {L1} and q_(001)_ {L2}. B) Detailed q range from A. C) Schematic illustration of the experiment based on free sedimentation and adsorption of LPX on the basal PEM. D) 1D GISAXS diffractogram of silicon wafer deposited LPX compared to si-(HA, CHI)_25_HA PEMs deposited after an adsorption period of 120 min in the in operando experiment. E) Detailed q range from D. F) Schematic illustration of the experiment based on free sedimentation and adsorption of LPX on the bare silicon wafer.

### 3.3. General Discussion

The presented experiments demonstrate that LPX composed of the lipid mixture OH4/DOPE loaded with DNA to a N/P ratio of 4 can successfully adsorb to HA/CHI PEMs (*in-situ* experiment, Figure 5A/B) and also be embedded in the PEM structure (Figure 4D/E). The PEM components HA and CHI are both weak polyelectrolytes which are loosely associated in a swollen PEM with high water content (Almodóvar et al., 2011) and, consequently, produce a surface coating with gel-like structure. The PEM-adsorbed LPXs seem to have no preferred orientation in the coating, indicated by the GISAXS pattern showing the Bragg reflection homogenously distributed along the Scherrer ring (ring labelled q_(001)_ in Figure 4C). When in contact with the HA/CHI PEMs, the repeat distance *d* of the lamellar LPX structure decreases by ≈7-10 Å compared to the LPX in dispersion, at least for a certain fraction of deposited PEMs. This observation needs to be discussed in more detail. The *d* value is the sum of the lipid bilayer thickness *d*_BL_ and the interlamellar water layer thickness *d*_W_ which also contains the DNA (d = *d*_BL_ *_+_ d*_W_). Hence, a smaller *d* value can result from a decreased *d*_BL_ and/or *d*_W_. An effect of the PEM adsorption on *d*_BL_ can be excluded as dominating factor for the increase since OH4 and DOPE as well as the mixtures are already in the liquid-crystalline phase state (high amount of *gauche*-conformers) (Janich et al., 2014) (Janich et al., 2015). Consequently, a reduced *d*_W_ must be responsible for the observed phenomenon. Hypothetically, the loss of the nucleic acid cargo DNA from the LPX could explain a decrease in *d*_W_, although the diameter of the DNA helix is 20 Å (https://doi.org/10.1021/ma00194a048). This hypothesis can be withdrawn by the fact that experiments with fluorescently-labelled LPX (lipid and DNA co-labelling) embedded in HA/CHI PEM indicate DNA-loaded LPX, and also LPX transfection functionality after PEM embedding was maintained (Krabbes et al., 2025a) (Husteden et al., 2020). Further, HA has a lower charge density than DNA (Delač Marion et al., 2015) (Vuletić et al., 2010) and consequently should be unable to disintegrate the complex of the cationic lipid bilayers with DNA. We already demonstrated that chondroitin sulphate, a polysaccharide with higher charge density compared with hyaluronic acid (Almodóvar et al., 2011), could not disintegrate OH4/DOPE NP4 LPX (Janich et al., 2015). Only heparin is able to disintegrate OH4/DOPE NP4 LPX do to the extraordinarily high charge density.

This brings us to the most likely hypothesis, a decreased *d*_W_ caused by a dehydration of the DNA/water layer between the lipid bilayers. But which phenomenon can explain this loss of water in the structure?

In earlier research we showed that HA/CHI PEMs terminated with HA (a thinner PEM with the sequence [HA,CHI]_5_HA was used) provide a negative surface zeta potential at pH >3 (Husteden et al., 2020), a prerequisite for effective deposition of OH4/DOPE NP4 LPX with a positive zeta potential. In this preliminary work, we also postulated a deformation of PEM-deposited LPX, as indicated by a discrepancy between the LPX layer thickness inside the PEM structure determined via ellipsometry and the OH4/DOPE NP4 LPX size in solution (Husteden et al., 2020). In the present work we also examine the embedding of OH4/DOPE NP4 LPX in HA/CHI PEMs, and although the PEM layer has a higher thickness and is produced with a different buffer, we can also assume that the positively charged LPX interact with a negatively charged surface (HA-terminated PEM). The reduced *d* value supports the hypothesis of LPX deformation. The question arises which forces induce the effect. Is it based on electrostatic effects and mechanical forces, or does the difference of the physicochemical environment in PEMs result in the observed effects. Hence it was demonstrated that the dielectric properties and polarity of the environment in a PEM can be different from the bulk (Neff et al., 2006) (Durstock and Rubner, 2001) (Tedeschi et al., 2001) (Schönhoff et al., 2007). Also the internal pKa value of the polyelectrolytes in the multilayer system is affected (Sui and Schlenoff, 2004) (Rmaile and Schlenoff, 2002). Consequently, also local pH shifts in the PEM can be discussed, also affecting LPX structure as discussed above.

The effect of dehydration on the LPX structure seems to be caused by the contact of LPX to the HA/CHI PEM, furthermore the cover layer seems not necessary for this transition, because we observed the phenomenon also in the *in-situ* experiment (Figure 5), where no cover layer was applied. Nevertheless, the question is unresolved, whether LPX, which are directly in contact with the PEM, is deeply absorbed into the PEM, or the PEM-buffer interface has a different physicochemical environment compared to the bulk which dominates the LPX environment and explains the appearance of only LPX with the lower *d* value (L2-LPX) at the lowest examined LPX loading concentration (see Figure 2D/E and Table 2). Hence this effect can be saturated when higher amounts of LPX are deposited. In that case coexistence of the dehydrated L2-LPX and the fully hydrated L1-LPX species was observed (in presence and absence of the cover layer, see Figure 4/5). We assume that the fraction of embedded LPX in direct close contact with the PEM components is dehydrated, while a second fraction of LPX is prevented from contact with the PEM components due to interactions with other LPX or the bulk (see illustrations Figure 6).

**Figure 6.**
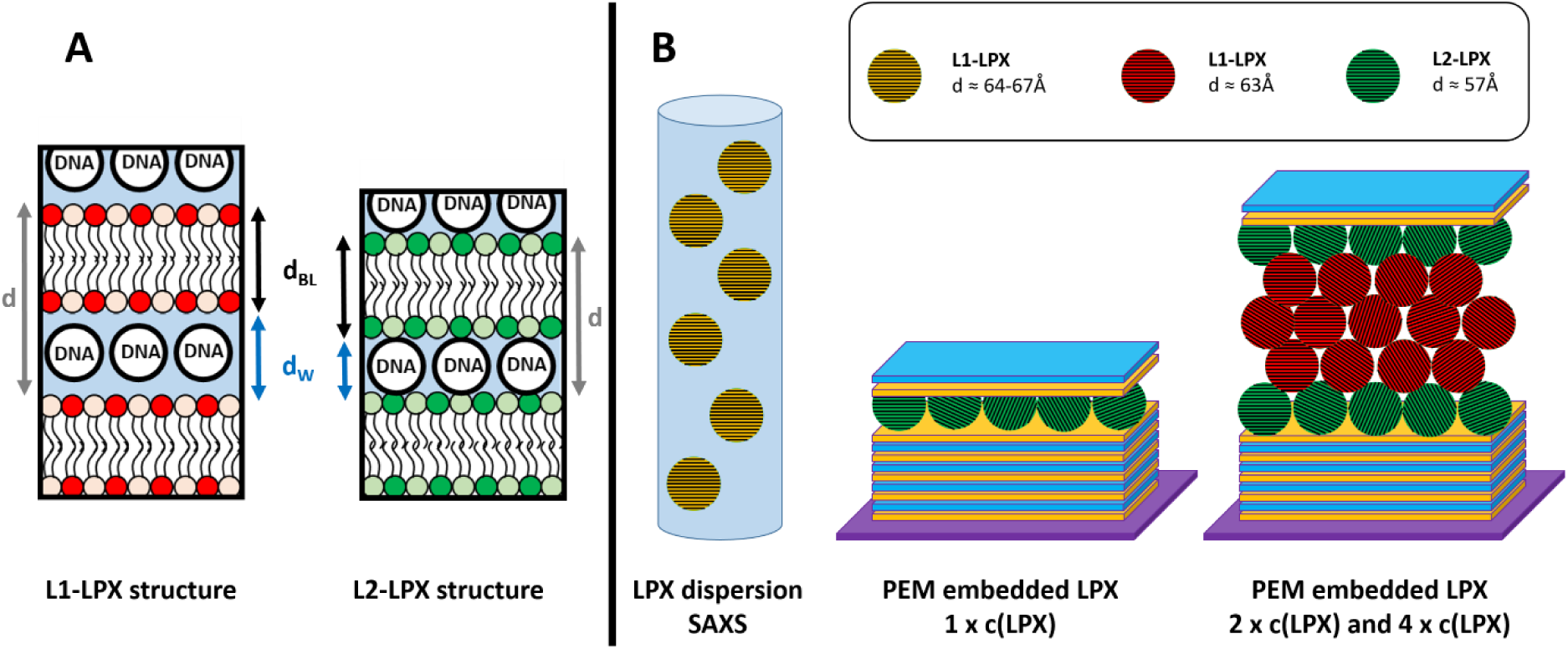
A) Schematic illustration of the proposed differences of the two lamellar LPX structures. B) Schematic illustration of the proposed localization of the different lamellar LPX structures in the samples.

The dehydration of LPX can be of biological relevance. For RNA-loaded LNPs it is proposed that the internal water content is connected with RNA degradation and consequently storage instabilities (Feng et al., 2025) (Schoenmaker et al., 2021). In previous work we also could demonstrate stabilizing effects of pluronics, polyethylene glycol-polypropylene glycol-polyethylene glycol block polymers, on siRNA loaded lipofectamine RNAiMAX nanoparticles (Mahmoud et al., 2023). It may be possible, that also here osmotic dehydration effects improved storage stability. In conclusion, we hypothesize that the controlled embedding of lipoplexes in PEM with dehydrating effects can be used to produce nucleic acid delivery systems with improved storage stability.

## 4. Conclusion

In this work we performed structural investigations of LPX, which were embedded in polyelectrolyte multilayers, via GISAXS. The technique allows to determine the effect of LPX immobilization in PEM on the internal structure of these nucleic acid/lipid nanoparticles. The studies show that the interaction of LPX with HA/CHI PEM is associated with changes in the repeat distance of the lamellar mesoscopic substructure of LPX. We propose that the interaction of lamellar LPX with PEMs result in a dehydrating effect on the lipid nanoparticles.

## Acknowledgement

We acknowledge Federal Ministry of Education and Research (BMBF) of Germany for funding Eiger2 1M detector via ErUM Pro 05K19FK2 (Murphy, CAU Kiel). Financial support for this research was provided by the Deutsche Forschungsgemeinschaft (DFG, German Research Foundation) – project number 396823779 (MK and CW). We acknowledge DESY (Hamburg, Germany), a member of the Helmholtz Association HGF, for the provision of experimental facilities. Parts of this research were carried out at PETRA III, beamline P08. Beamtime was allocated for proposal R-20240669. We thank the Biomaterials department of MPIKG for access to their X-ray infrastructure and Daniel Werner for help with the SAXS measurements. Parts of Figure 2 are provided by Servier Medical Art (https://smart.servier.com/), licensed under CC BY 4.0 (https://creativecommons.org/licenses/by/4.0/).

