## Supplemental Material for "Intrinsic structure of lipoplexes embedded in polyelectrolyte multilayers"

**Content:**

1. **SAXS results**
2. **Samples of GISAXS at the 3 different angles of incidence**
3. **Stored samples**
4. **Position scans of the different GISAXS samples**
5. **Gel for quantification of the DNA loading**
6. **XRR measurements**

### 1. SAXS results

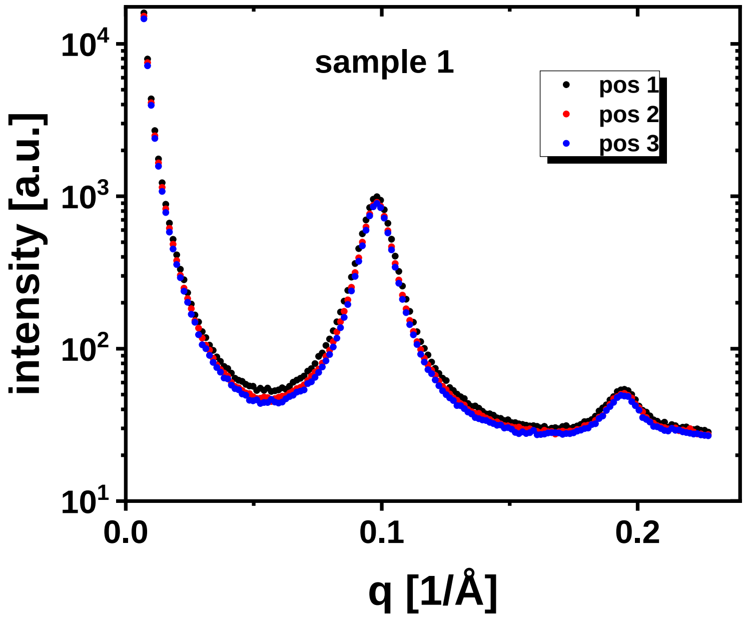


**Figure S1:** SAXS pattern of OH4/DOPE NP4 LPX sample 1 in acetate buffer pH 5.5 without baseline correction at 3 different positions of the capillary.

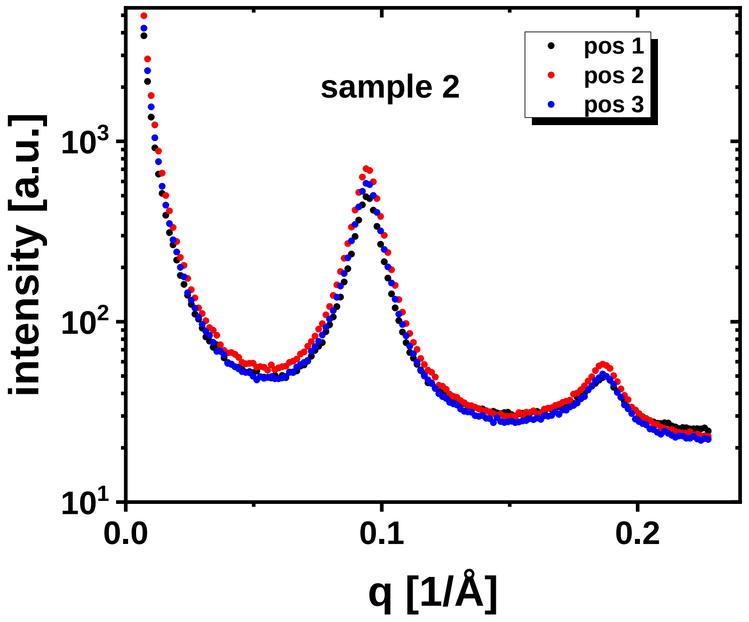


**Figure S2:** SAXS pattern of OH4/DOPE NP4 LPX sample 2 in acetate buffer pH 5.5 without baseline correction at 3 different positions of the capillary.

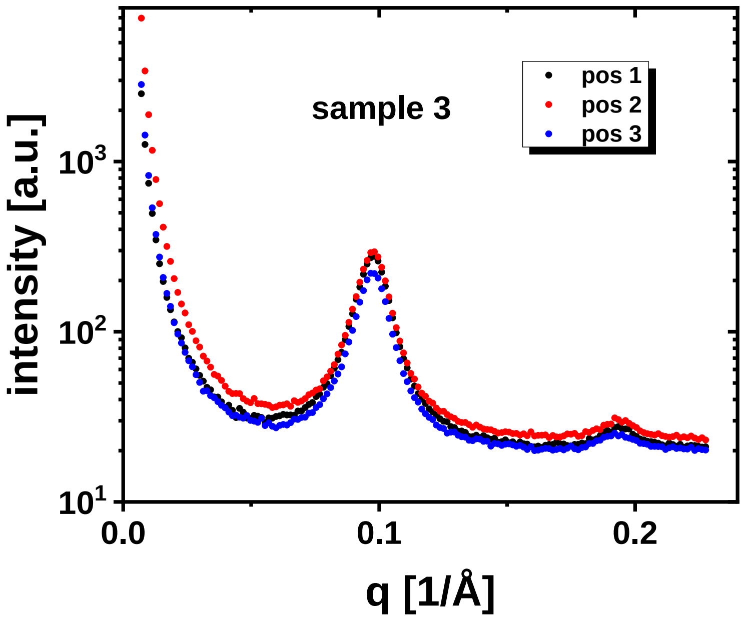


**Figure S3:** SAXS pattern of OH4/DOPE NP4 LPX sample 3 in acetate buffer pH 5.5 without baseline correction at 3 different positions of the capillary.

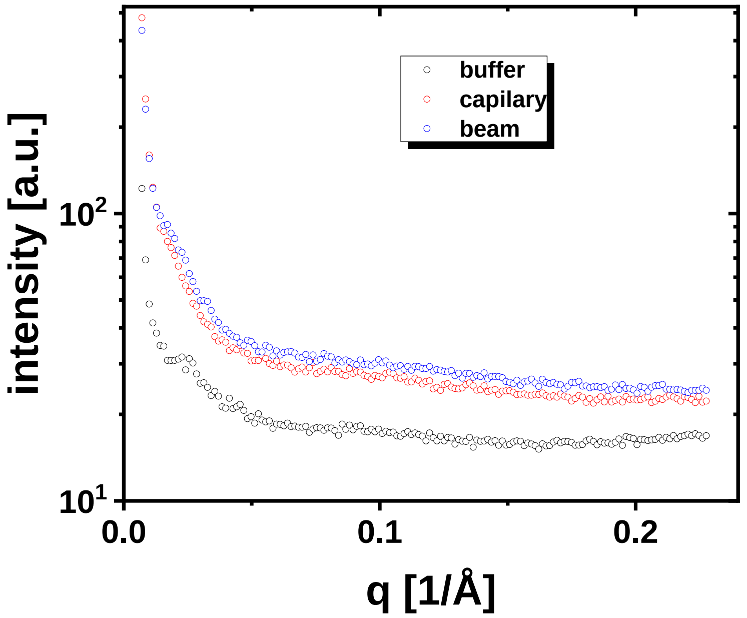


**Figure S4:** SAXS pattern background of the capillary filled with buffer (buffer), the empty capillary (capillary) and the empty beam path (beam).

### 2. Samples of GISAXS at the 3 different angles of incidence

A clear Scherer rings were observed in the GISAXS signal. Nevertheless, in some samples a ”ghost ring” occurs from the same Bragg reflection of the deposited layer under the total reflection of the incident beam from the substrate, when the incident angle is lower than the critical angle of the substrate. At 18 keV, the measurement at 0.07° incidence meets this condition, as the critical angle of the air-silicon interface is about 0.095°. The dominant Bragg reflection occurs when the incident beam is directly diffracted by the lamellar structure in the deposited layer, exhibiting the strongest Scherrer ring. A secondary Bragg reflection occurs when the reflected beam from the silicon substrate surface re-enters the deposited layer and gets diffracted, exhibiting a weaker “ghost ring”. The ghost ring is weaker because the reflected beam from the wafer surface has been attenuated by the deposited layer once. Since the primary beams of the two diffraction events are the incident beam and the reflected beam from the substrate, their orientation differs in the vertical direction by two times the incident angle. Accordingly, the main diffraction ring and the ghost ring are offset vertically by two times the incident angle. At higher incident angles where the reflection from the wafer surface is much weaker, the ghost ring is negligible.

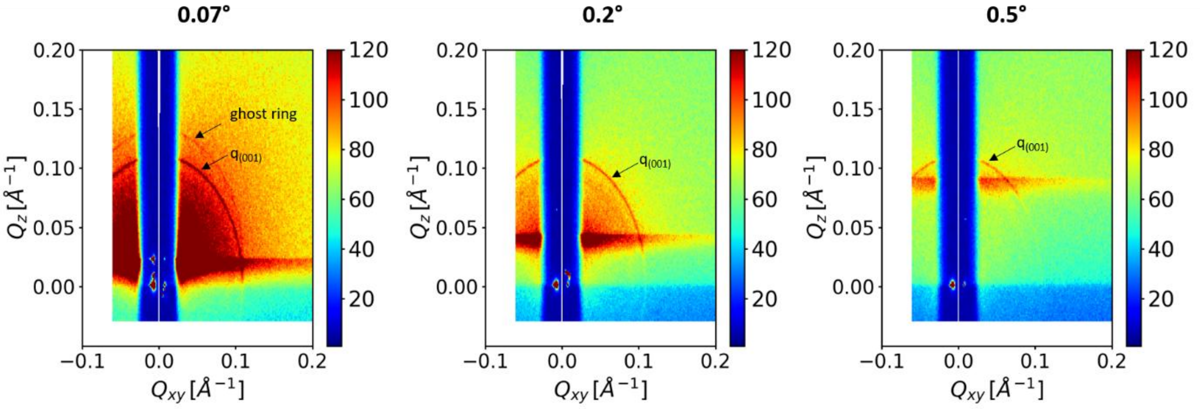


**Figure S5:** Detector image of the GISAXS signal of the coating sequence PEM-LPX-HA-CHI at different incidence angles.

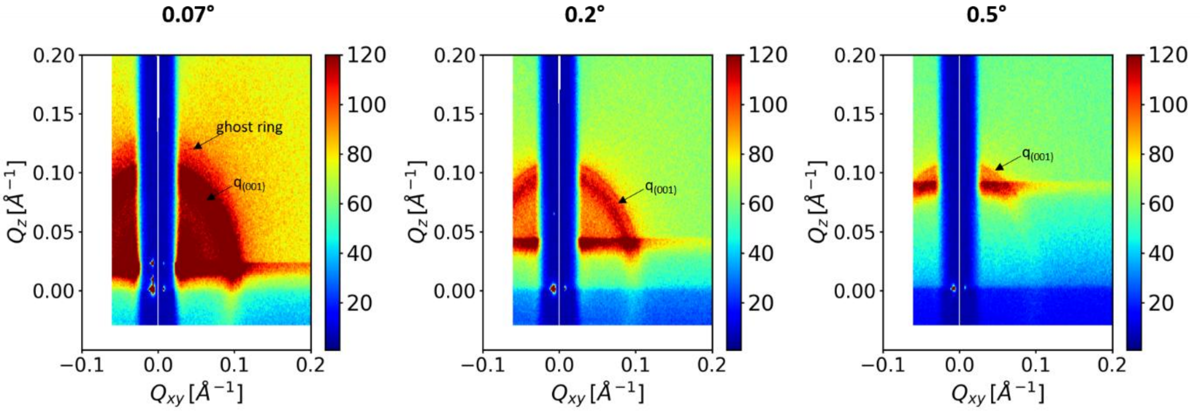


**Figure S6:** Detector image of the GISAXS signal of the coating sequence PEM-LPX-HA-CHI 4 x c(LPX) at different incidence angles.

### 3. Stored samples

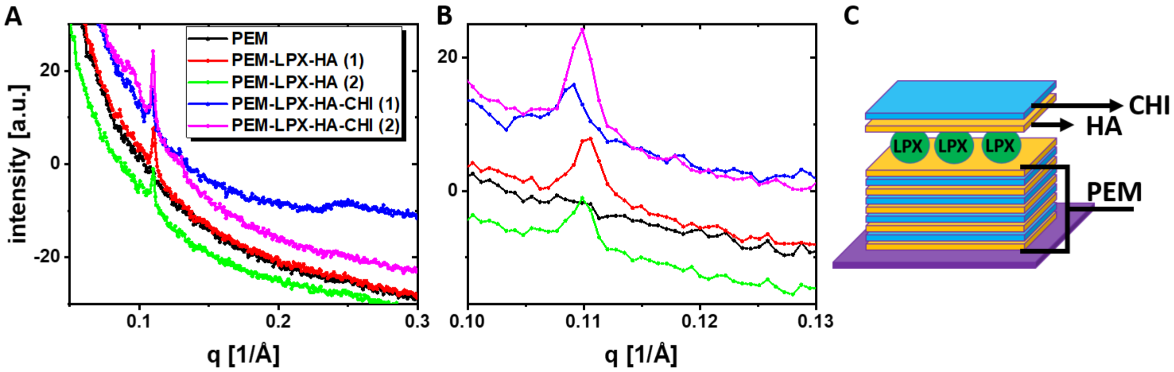


**Figure S7**: GISAXS experiments of stored samples at different processing levels. A) 1D GISAXS diffractogram of stored PEMs at different processing steps (depicted in C). (1) and (2) label two independent samples with the same structure. GISAXS diffractogram of the samples at 4 different positions are shown in the supporting information (Figure S8-11) to demonstrate homogeneity of the samples. B) Detailed q range from A.

**Table S1** LPX Bragg peak position obtained from GISAXS measurement of OH4/DOPE NP4 LPX on or embedded in PEMs as mean of 4 different positions

| | lamellar structure 1 $q_{(001)}$\{L1\} [d] | lamellar structure 2 $q_{(001)}$\{L2\} [d] | LPX loading dispersions: concentration | coverlayer on top of LPX |
|---|---|---|---|---|
| stored samples | | | | |
| PEM-LPX-HA (1) | - | 0.1101 1/Å [57.0 Å] | 4.33 ng/µL | HA |
| PEM-LPX-HA (2) | - | 0.1097 1/Å [57.3 Å] | 4.33 ng/µL | HA |
| PEM-LPX-HA-CHI (1) | - | 0.1089 1/Å [57.7 Å] | 4.33 ng/µL | HA/CHI |
| PEM-LPX-HA-CHI (2) | - | 0.1098 1/Å [57.2 Å] | 4.33 ng/µL | HA/CHI |

### 4. Position scans of the different GISAXS samples

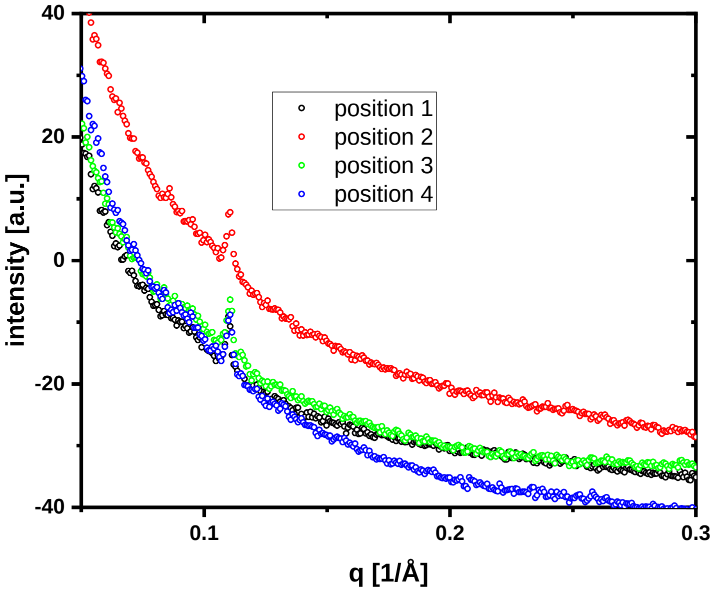


**Figure S8:** GISAXS diffraction patterns obtained by azimuthal integration of the 2D detector image at beam incidence of 0.07°of the stored PEM coated silica wafer with the PEM system PEM-LPX-HA (1) measured at 4 different positions.

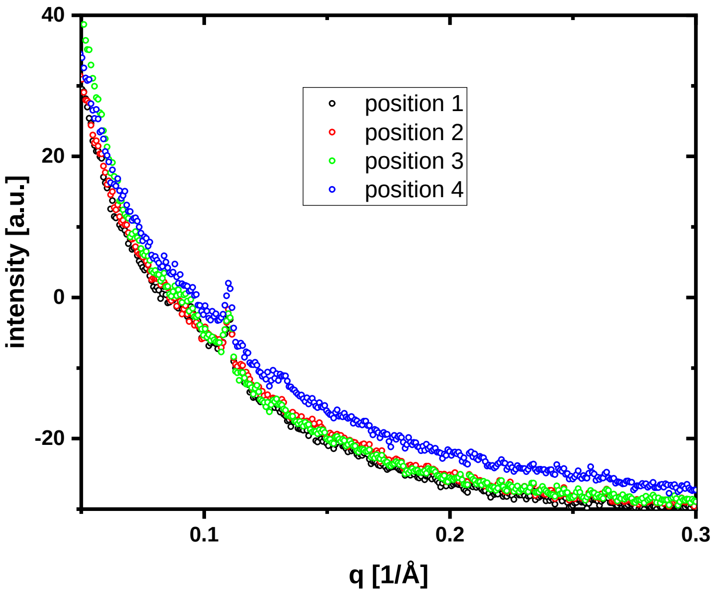


**Figure S9:** GISAXS diffraction patterns obtained by azimuthal integration of the 2D detector image at beam incidence of 0.07°of the stored PEM coated silica wafer with the PEM system PEM-LPX-HA (2) measured at 4 different positions.

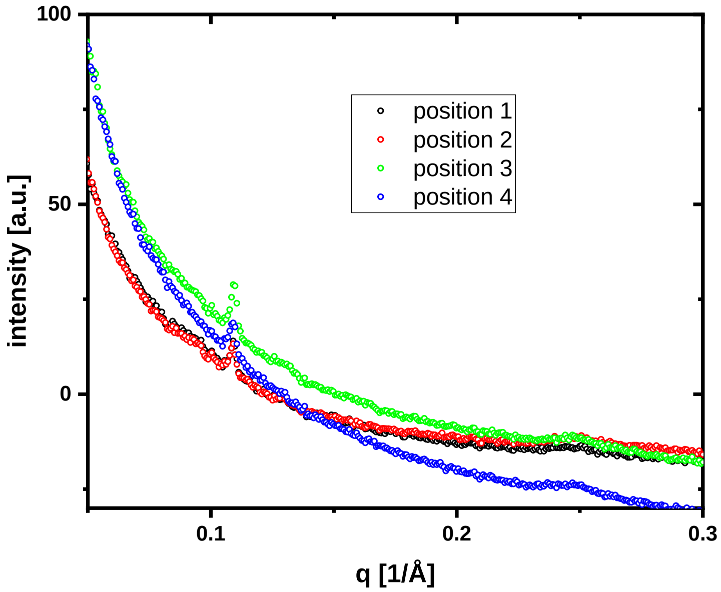


**Figure S10:** GISAXS diffraction patterns obtained by azimuthal integration of the 2D detector image at beam incidence of 0.07°of the stored PEM coated silica wafer with the PEM system PEM-LPX-HA-CHI (1) measured at 4 different positions.

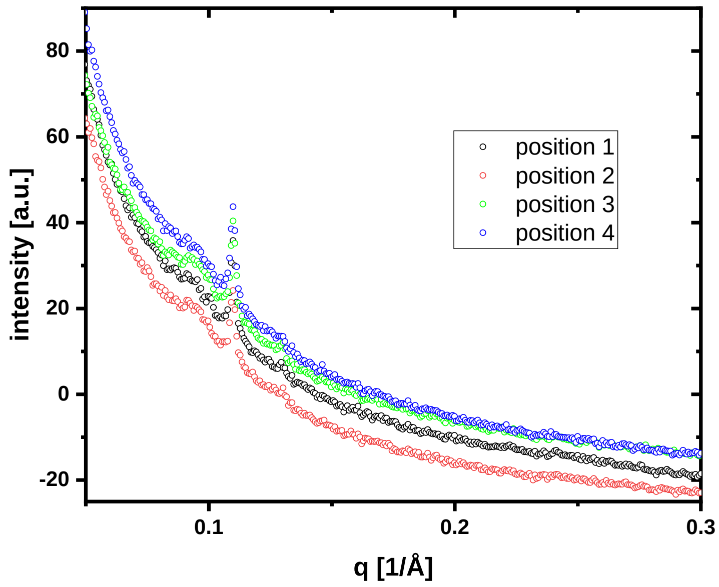


**Figure S11:** GISAXS diffraction patterns obtained by azimuthal integration of the 2D detector image at beam incidence of 0.07°of the stored PEM coated silica wafer with the PEM system PEM-LPX-HA-CHI (2) measured at 4 different positions.

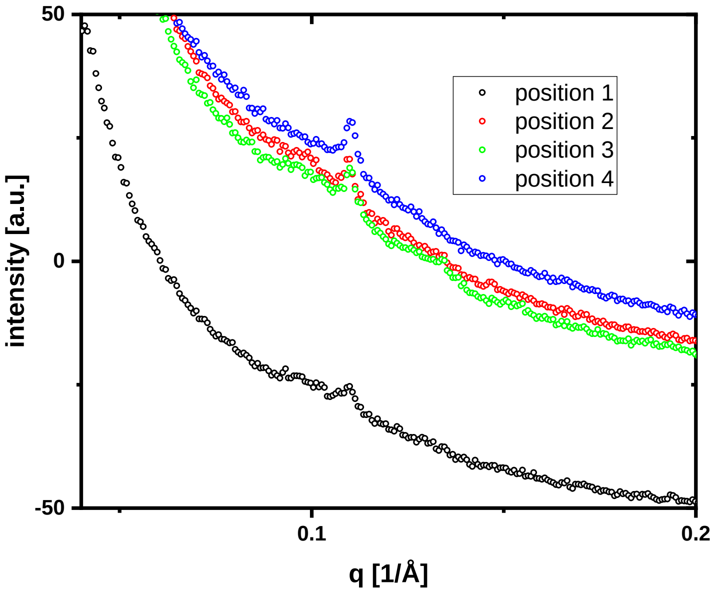


**Figure S12:** GISAXS diffraction patterns obtained by azimuthal integration of the 2D detector image at beam incidence of 0.07°of the silica wafer with the PEM system freshly coated with the PEM-LPX-HA-CHI coating with the 1 x c(LPX) loading quantity measured at 4 different positions.

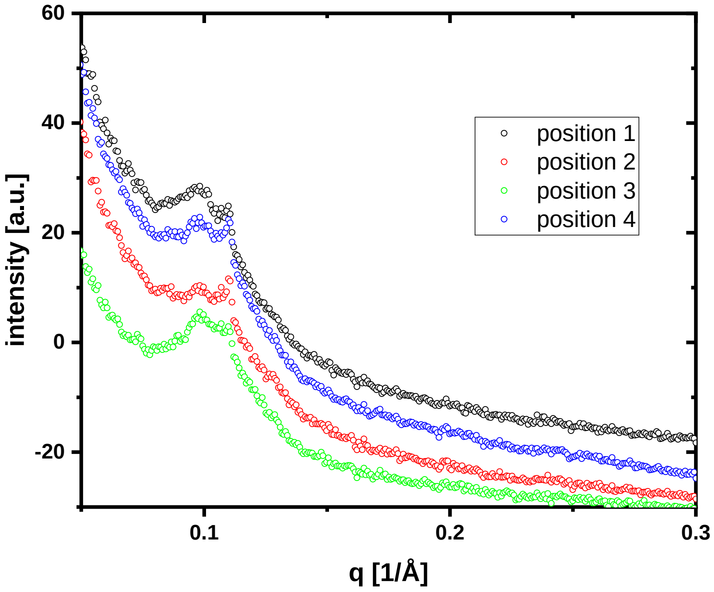


**Figure S13:** GISAXS diffraction patterns obtained by azimuthal integration of the 2D detector image at beam incidence of 0.07°of the silica wafer with the PEM system freshly coated with the PEM-LPX-HA-CHI coating with the 2 x c(LPX) loading quantity measured at 4 different positions.

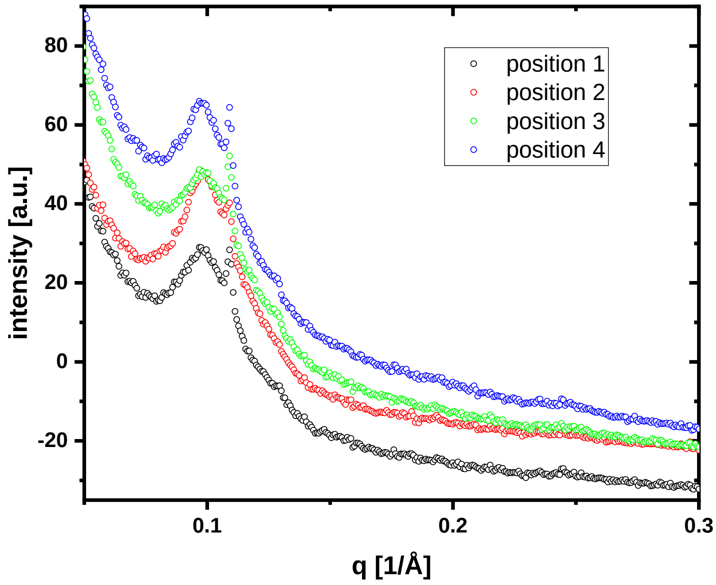


**Figure S14:** GISAXS diffraction patterns obtained by azimuthal integration of the 2D detector image at beam incidence of 0.07°of the silica wafer with the PEM system freshly coated with the PEM-LPX-HA-CHI coating with the 4 x c(LPX) loading quantity measured at 4 different positions.

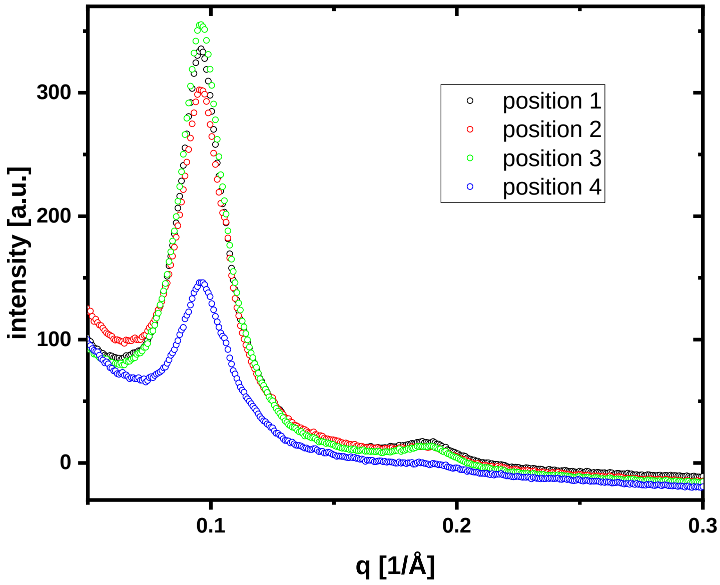


**Figure S15:** GISAXS diffraction patterns obtained by azimuthal integration of the 2D detector image at beam incidence of 0.07°of the silica wafer with the PEM system freshly coated with the PEM-LPX-HA-CHI coating with the 8 x c(LPX) loading quantity measured at 4 different positions.

### 5. Gel for quantification of the DNA loading

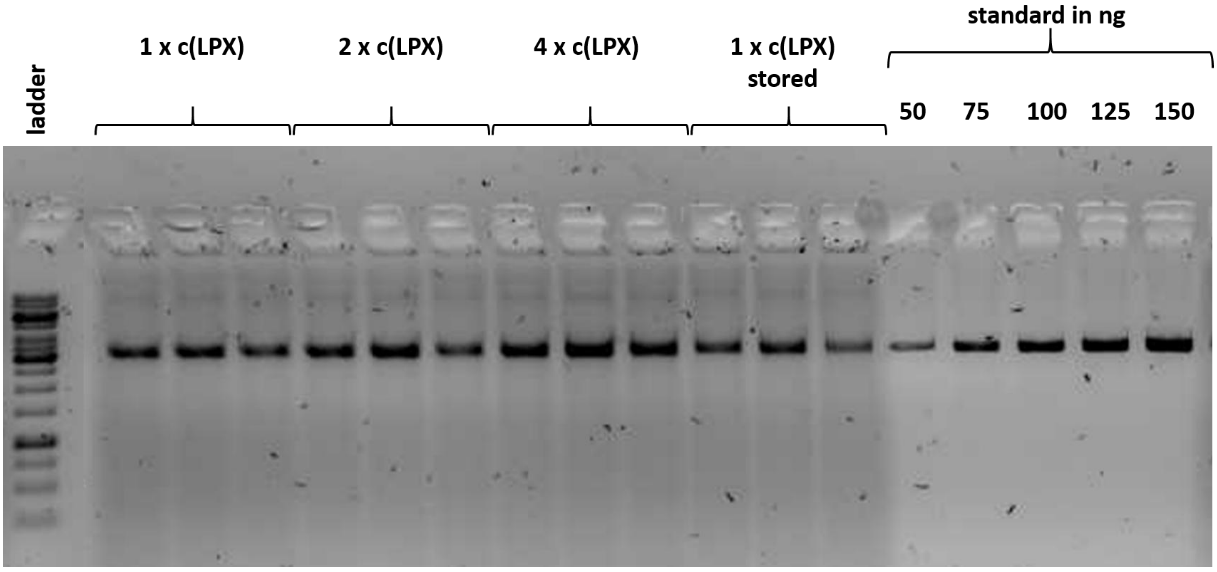


Figure S16: Gel electrophoresis of the DNA quantification procedure described in chapter 2.10 “Quantification of the DNA loading” of the main article.

### 6. Gel for quantification of the DNA loading

The XRR data comparing the base PEM (HA, CHI)$_{25}$HA with the LPX-loaded PEM of the sequence (HA, CHI)$_{25}$HA-LPX-HA-CHI show no pronounced difference between both curves (Figure 4G). The curves show no clear Kiessig fringe or Bragg reflections. This can be caused by a low contrast in electron density of the components or by the absence of any ordering in the direction perpendicular to the surface. Further it has to be noted, that the thickness of the swollen PEM layer is larger than a µm, so that the fringes associated with the whole layer will not be resolved by the XRR experiment. The small feature observed for the PEM sample at ≈1.7 1/Å may be caused by the silicon oxide layer of the silicon wafer.

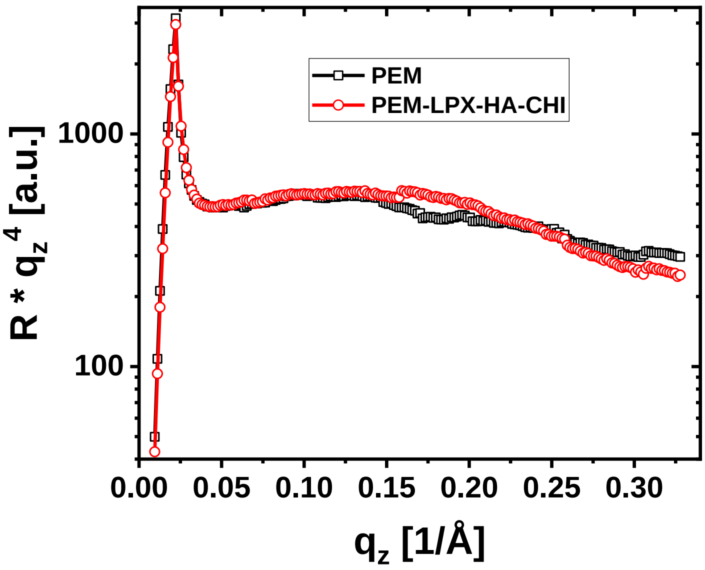


Figure S17: Reflectivity experiments of samples of the base PEM (HA, CHI)$_{25}$HA and the LPX loaded PEM of the sequence (HA, CHI)$_{25}$HA-LPX-HA-CHI in buffer.
